# Experimental design impacts the vector competence of *Ornithodoros* ticks for African swine fever virus: a meta-analysis of published evaluations

**DOI:** 10.1101/2023.11.10.566648

**Authors:** Jennifer Bernard, Sara Madeira, Joachim Otte, Fernando Boinas, Marie-Frédérique Le Potier, Laurence Vial, Hélène Jourdan-Pineau

**Affiliations:** CIRAD, UMR ASTRE, F-34398 Montpellier, France. ASTRE, Univ Montpellier, CIRAD, INRAE, Montpellier, France; Unité Virologie et Immunologie Porcines, Laboratoire de Ploufragan-Plouzané, Agence Nationale de Sécurité Sanitaire (Anses), Univ Bretagne Loire, BP53, 22440 Ploufragan, France; Faculty of Veterinary Medicine, University of Lisbon, Avenida da Universidade Técnica, 1300-477 Lisbon, Portugal CIISA—Center of Interdisciplinary Research in Animal Health, 1300-477 Lisbon, Portugal Associate Laboratory for Animal and Veterinary Sciences (AL4AnimalS), 1300-477 Lisbon, Portugal; Berkeley Economic Advising and Research. 1442A Walnut Street, Berkeley, CA, 94705 USA

**Keywords:** *Ornithodoros* soft ticks, African swine fever virus, vector competence, experimental methodology, meta-analysis

## Abstract

African Swine Fever (ASF) is a highly economically devastating viral disease for swine. Soft ticks of the genus *Ornithodoros* are involved in its epidemiology, as vectors and natural reservoirs of African Swine Fever Virus (ASFV). The vector competence of *Ornithodoros* ticks for ASFV has been mainly studied by mimicking natural tick-to-pig transmission pathways through experimental infections in the laboratory. By reviewing the original research studies dating back to the 1960s on the vector competence of *Ornithodoros* for ASFV, we estimated the vector competence of 10 tick species in association with 38 viral strains resulting in 51 tick-virus associations. This assessment emphasized the extensive range of protocol designs employed and their impact on the success of tick infection with ASFV. Our results call for standardised procedures in vector competence experiments to facilitate further investigation and reduce potential experimental bias. In particular, we recommend the use of late nymphs or adult ticks from a laboratory colony to achieve efficient infection rates. In addition, viral inoculation should be carried out by blood meal rather than by injection, and preference should be given to high titre blood. Finally, detection of viral DNA should be performed 2 months after inoculation to distinguish between successful replication and residual virus in the tick.

## Introduction

African swine fever (ASF) ranks among the most damaging viral diseases affecting swine farming. The etiologic agent is a large DNA virus belonging to the *Asfaviridae* family (Dixon et al., 2013). African Swine Fever Virus (ASFV) causes haemorrhagic fever in susceptible infected swine, with mortality rates of up to 100% in naïve domestic pigs. In Africa, wild suids (warthogs, and possibly bushpigs and giant forest hogs) are natural vertebrate reservoirs that are resistant to the disease (Jori et al., 2013). In Europe, wild boars are susceptible to ASFV and exhibit clinical symptoms and mortality rates comparable to those of domestic pigs. In certain countries, soft ticks of the genus *Ornithodoros* act as vectors or natural reservoirs of ASFV. The virus can persist in ticks for several years, potentially leading to epizootic outbreaks in pig farms that have become colonized by ticks, even with the implementation of ASF control measures (i.e., pig slaughtering, disinfection, and quarantine) (Boinas et al., 2011; Jori et al., 2013). Once ASFV is introduced into a new free area where the presence of *Ornithodoros* ticks is highly suspected, as it occurred in the Caucasus in 2007 or in the Caribbean in 2021, the potential for long-term persistence of ASFV in tick reservoirs and ASFV transmission via tick bites raises concern (Beltrán-Alcrudo et al., 2008; EFSA et al., 2021). *Ornithodoros* soft ticks feed not only on wild suids but on a large variety of vertebrate hosts (aardvark, porcupine, chicken tortoise…). However, the presence of ASFV in these non-suid hosts has never been detected.

Vector competence generally refers to the vector’s capacity to be infected by a pathogen, to maintain and multiply this pathogen, and finally to transmit it to a new susceptible host (Gillespie et al., 2004). The biological steps taking place within the arthropod vector can be broken down into four stages: (i) the ingestion of an infected blood meal, (ii) the infection of the vector upon crossing of its intestinal midgut by the pathogen, (iii) the persistence and multiplication of the pathogen within the vector, and (iv) the dissemination of the pathogen inside the vector by crossing organic barriers and ultimately reaching organs implicated in the transmission to vertebrate hosts, typically the salivary glands and coxal glands. Sexual transmission of pathogens between males and females of vectors as well as transovarial transmission from female vectors to their progeny can occur through the infection of reproductive organs. Although not strictly included in the formal definition of the vector competence, these pathways of transmission will also be considered in this review.

All these biological steps are observed in the interaction between ASFV and *Ornithodoros* tick although results reported varied greatly between studies. In addition to historical demonstration of tick vector competence for ASFV in African *O. moubata sensu lato* and Iberian *O. erraticus*, several other *Ornithodoros* tick species succeed to maintain or transmit ASFV under experimental conditions (Mellor & Wilkinson, 1985; Endris et al., 1987, 1991; Hess et al., 1987). This suggests that most *Ornithodoros* ticks might be able to maintain and transmit ASFV (EFSA Panel on Animal Health and Welfare., 2010a). Based on available results, some researchers have proposed to predict the global role of each tick species as a vector and reservoir for ASFV (Kleiboeker & Scoles, 2001; EFSA Panel on Animal Health and Welfare., 2010b; a, 2014; Burrage, 2013). However, these comprehensive reports do not account for the evident disparities in vector competence results, which likely rely on intricate tick-virus adaptations as proposed by Pereira de Oliveira (Pereira de Oliveira et al., 2019; Pereira De Oliveira et al., 2020) and other yet-to-be-identified biological and experimental factors. No paper has considered the substantial diversity of techniques employed to test vector competence except for a study on *O. erraticus* infected with two Portuguese ASFV strains (Ribeiro et al., 2015). Nonetheless, the importance of experimental methodology has been demonstrated in other models like mosquitoes with Rift Valley fever, Dengue, Zika or Japanese Encephalitis viruses (Azar, 2019; Auerswald et al., 2021; Drouin et al., 2022). We hypothesize that the experimental design, including the choice of biological materials and the methods used to infect ticks and detect the virus, is an important factor in explaining variation in vector competence, in addition to the influence of tick species, ASFV strains and their combinations.

To investigate this hypothesis, a systematic review of studies published up to 2024 was conducted, examining the vector competence of *Ornithodoros* ticks for ASFV. The present paper focuses on the diverse experimental factors that may impact vector competence and determines the overall vector competence of the various associations between *Ornithodoros* tick and ASFV. We provide recommendations on how experimental trials can be refined to minimize biases and obtain more comparable and reliable results on *Ornithodoros* vector competence for ASFV.

## Material and Methods

### Collection of bibliographical resources

The first objective of the study was to identify the bibliographic resources related to vector competence of *Ornithodoros* ticks for ASFV. For practical issues, we decided to consider only resources published in the English language.. This was achieved following the Preferred Reporting Items for Systematic Reviews and Meta-Analyses (PRISMA) guidelines (Annex 1). Our bibliographic review was conducted until July 2022, using three different bibliographic databases: EThOS (http://ethos.bl.uk/Home.do), PubMed (http://www.ncbi.nlm.nih.gov/) and Scopus (http://www.scopus.com/). Queries were built under the keywords “African swine fever” AND “tick”. The topics not related with tick competence (e.g., immunology, genetic, vaccinology, epidemiology) were excluded during the screening phase. Then, during the eligibility phase, studies were excluded if quantitative reliable data could not be extracted from publications. Additionally, reviews lacking original vector competence results and offering merely descriptive information were excluded. In addition to scientific articles published in peer-reviewed journals, PhD manuscripts were also selected.. All duplicated results were removed to avoid biases in the analyses. Finally, a review of the citations in the included references was carried out in order to identify further relevant studies. The final list of included references that were used to extract all data in soft tick vector competence for ASFV are shown in Table 1.

**Table 1.**
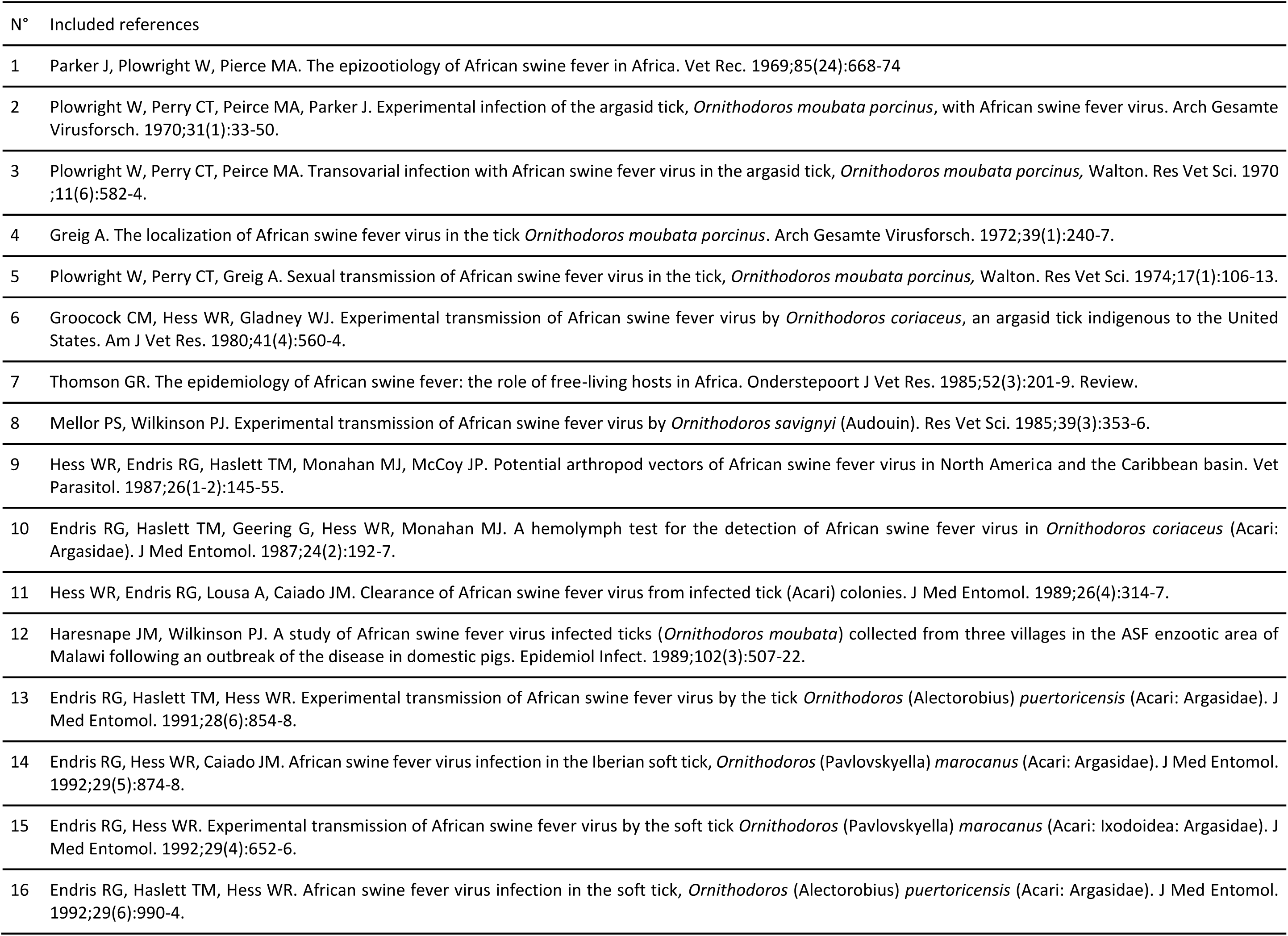
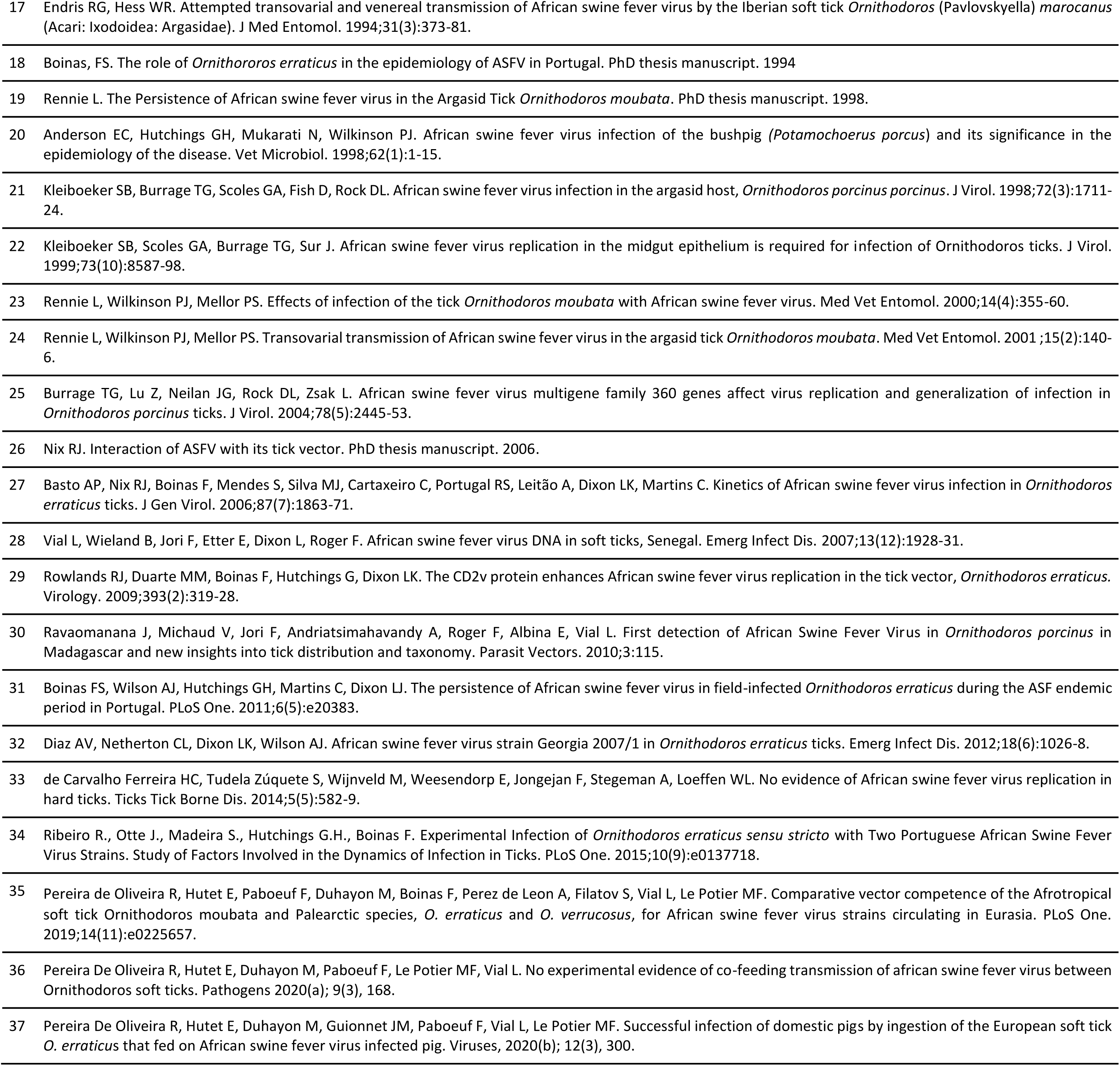
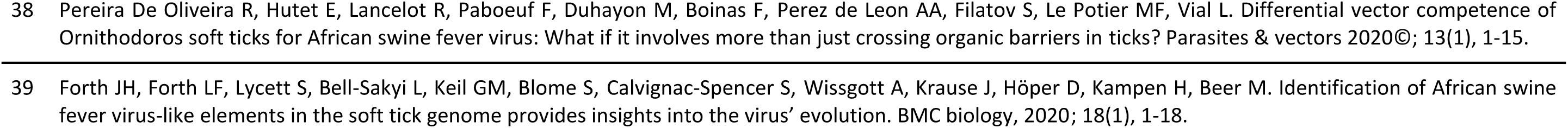
Included references providing original results on the vector competence of *Ornithodoros* ticks for ASFV, with number attributed to each paper by year of publication.

### Extraction of data about tick competence from bibliographical resources

General bibliometric information on authors and years of publication was collected to describe the general interest in tick vector competence for ASFV and to identify the community of scientific experts. We considered the experts as the persons in first and last position of the included references.

Information on tested *Ornithodoros* tick-ASFV associations was systematically reported. Regarding virus, we recorded the strain, the genotype (on the basis of p72 gene sequences, Qu *et al*. 2022), the geographical origin, and the host (suids or ticks) from which the ASF virus was isolated. The original names of some ASFV strains were retrieved by crossing information from the literature, according to known authors’ collaborations, or by directly contacting the authors. When the determination of strains remained impossible, the strain was identified by indicating the country where the ASFV strain was isolated, the year of the study, and a letter if duplicates existed. One exception was a strain used to infect *O. turicata* (Hess et al., 1987) without any indication of the country of origin which was named “Unknown”. Six recombinant ASFV strains, genetically modified and obtained from two natural strains (Burrage et al., 2004; Nix, 2006), were excluded from this review since they cannot infect ticks in nature. Genotypes were not available for some strains of ASFV, but they could be inferred based on their geographical origin and ASFV genotype distribution published by Lubisi et al. (2005).

Concerning ticks, the species, their geographical origin, their colony origin and type were reported. When ticks were reared for less than five years in the laboratory, they were considered comparable to ticks sampled from the field due to the limited number of generations occurring during this period. Soft tick systematics has greatly changed during the last decades. Tick species identification was updated in accordance with the revision of valid tick names by Guglielmone (Guglielmone et al., 2010) and the mapping of their known distribution (Vial, 2009). The more recent taxonomy study of southern African *Ornithodoros* ticks (Bakkes et al., 2018) could not be used, due insufficiently precise location to attribute a species name among the *O. moubata* complex of species. Finally, when determining tick geographical origin, 6 studies involving 4 different tick species (*O. coriaceaus, O. parkeri, O. moubata*, and *O. porcinus*) did not report this information but since *O. parkeri* and *O. coriaceus* are known to be limited to North America, especially the United-States (De La Fuente et al., 2008), they were assigned to this country.

Authors used various measurements to assess the tick vector competence for ASFV. Some authors reported numbers of ticks positively detected with ASFV, while others scrutinized the virus titres in ticks and tick organs or monitored ASFV kinetics for several weeks or months, providing more comprehensive information on the virus persistence within its tick reservoir. Even if ASFV detection in tick transmission organs (salivary or coxal glands, gonads) suggest possible successful transmission to pigs or to other ticks, this alone is insufficient to confirm such transmission pathways. Conversely, the detection of ASFV in transmission secretions (e.g., saliva or coxal fluids) is stronger evidence, as the virus is already excreted in this case. In these transmission experiments, clinical signs observed in pigs following tick bites and detection of ASFV in pig blood provide definitive confirmation of the ability of ticks to transmit the disease. Similarly, detection of ASFV in tick sexual mates, offspring confirms sexual and vertical transmission. The transmission of virus between ticks via co-feeding has been described in hard ticks, but this phenomenon has not been demonstrated for ASFV in soft ticks (Pereira de Oliveira et al., 2020).

Based on these data, it was possible to evaluate three major components of the vector competence of the ticks: 1) the tick infection (revealed by the crossing of the tick midgut by ASFV), 2) the ability of ticks to transmit ASFV to pigs and 3) their ability to transmit ASFV to other ticks). Transmission to pig through tick bite or from ticks to ticks cannot occur without the previous infection of the tick, which was thus considered as the main step to be assessed. In addition, tick ability to transmit ASFV to pigs or other ticks are less frequently reported (N= 160 and N=54 over 900 experiments, respectively) likely because they are somewhat more difficult to assess than tick infection. Therefore, for each experiment, we computed a binary variable, the infection status, according to any measurements providing direct information on tick infection as well as indirect transmission measurements that imply previous tick infection (Figure 1). More precisely, if ASFV transmission to pigs or ASFV transmission from ticks to other ticks were reported in papers, the ticks were considered to be infected as the virus has necessarily crossed the tick organic barriers to reach transmission organs and be transmitted. If transmission failed or was not tested, the detection of ASFV in ticks was used as an indication of infection status. Obviously when ASFV was not detected, ticks were considered as not infected. The only exception was when the detection occurred on the same day as virus exposure, in which case no virus detection indicates a failure of experimental infection or a too low detection threshold.

**Figure 1.**
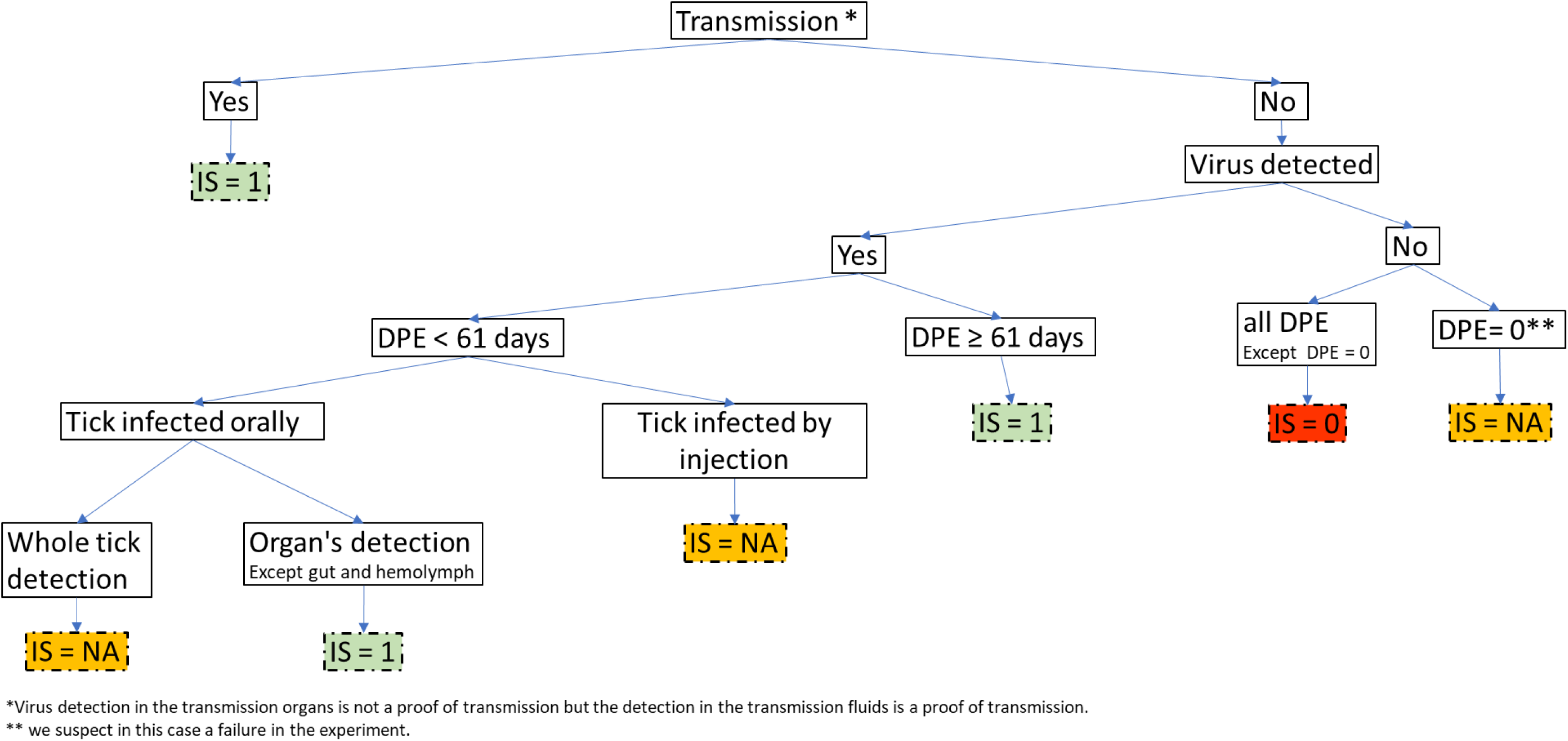
Criteria used to determine the infection status of ticks after their virus exposure, with three classes (IS: Infection Status; DPE: Date post exposure; NA: Not Applicable, 1: Infected, and 0: Not Infected).

Different methods have been used by authors to detect ASFV in ticks, namely ASFV isolation/titration, observation of ASFV particles, and viral DNA or RNA detection using multiple designs of PCR. In the included references, ASFV isolation/titration was based on haemadsorption (HAD), which is the ability of many ASFV strains to aggregate on red blood cells when they infect pig alveolar macrophages and create typical “rosettes” (Malmquist & Hay, 1960). . This technique is largely used by authors but may suffer from inhibition issues due to cytotoxic effect of tick components on pig macrophages and its achievement in at least 6 days is very time-consuming (Oura et al., 2013). Virus titration implies that the virus is able to replicate but does not necessarily confirm that it has crossed the midgut barrier. Similarly, detection of viral RNA by Reverse Transcription quantitative PCR also indicates active viral replication. Conversely the detection of viral DNA can be due to residual virus from blood meal and does not mean that the virus is still active. However, we considered that those methods brought equivalent results to assess infection status once the duration after exposure was taken into account.

Indeed, when ASFV was detected in ticks, a minimum duration of 61 days between tick exposure to ASFV and detection of ASFV in tick (hereafter DPE for days post exposure) was chosen to distinguish successful viral replication inside the tick from detection of residual virus after the infectious blood meal or virus inoculation (Figure 1). Indeed, the minimum duration for blood meal digestion and clearance of ingested blood in soft ticks is approximately 4 weeks (Rowlands et al., 2009). Furthermore, if tick infection is unsuccessful, ASFV can be detected during 4 weeks by isolation (Ribeiro *et al*., 2015) and after at least 49 days by qPCR (de Carvalho Ferreira et al., 2014; see also Bernard, 2015 for a review on ASFV kinetics inside soft ticks). If tick infection is successful, virus generalization in ticks takes about 30 to 56 days (Plowright et al., 1970; Kleiboeker et al., 1998; Basto et al., 2006; Burrage, 2013). Therefore, and as a precaution, if ASFV was tested and detected before 61 DPE, the infection status was considered unknown unless virus was detected in organs other than the midgut. In this case, this indicates that the virus has crossed the midgut barrier.

Inoculation by injection allows accurate assessment of the virus dose received by the tick. As it delivers the virus directly into the haemocoel, circumventing the midgut barrier, it offers valuable insights into the molecular compatibility between the virus and the tick. However, it is acknowledged that the injection can be traumatic for the tick, resulting high mortality and leads to distribution of the virus throughout the tick (Bonnet & Liu, 2012). In this case, as the detection of virus in the organs was doubtful, the infection status was recorded as unknown.

The experimental design used in the different studies was also reported, including: i) the exposure route for the tick infection, ii) the ASFV titres in the inoculum used to infect the ticks (infectious dose), iii) the development stage of the ticks tested in the study, iv) the DPE, v) the number of ticks tested in the study. In field studies, viral titres in the hosts were not necessarily reported but they were considered to be high (>10 ^6^ HAD50/mL of blood) considering the viraemia usually observed in ASF diseased pigs (Plowright et al., 1968). Tick stage was classified as “early” from larva to nymph 3 and as “late” from nymph 4 to adult.

### Statistical analysis of the tick infection status

A statistical analysis was carried out to identify factors potentially affecting the tick infection status. More precisely, we investigated if factors related to the nature of tick-virus association or the experimental design influenced the infectious status (see above for details) of the ticks in published experimental trials. The explanatory variables were: the ASFV strain and genotype, the tick species, the potential geographical co-occurrence of tick and virus as an indication of possible tick-virus adaptation (ASFV and ticks from the same country versus from different countries), the viral titre in infectious blood meal and the exposure route for tick infection, the tick stage, the tick colony type and the host from which the ASFV strain was sampled. Using a dataset where all ticks were adults, we also tested the influence of tick sex on infectious status (along with all other factors except tick stage). All statistical analyses were performed on R 4.2.2 (R Core Team, 2022). To assess which factors influenced the infection status, we applied a generalized mixed model with a binomial family. The tick species, virus genotype and virus strain were set as random effects to account for the fact that only a random fraction of soft tick species and virus genotypes and strains were sampled. All other factors were set as fixed effects. The significance of random effects was tested using a likelihood ratio test (Zuur et al., 2009). We then selected the best parsimonious models (with the remaining 6 fixed effects) based on the AIC (Akaïke Information Criterion). In this best model, the significance of fixed effects was tested using Chi² tests between models and post-hoc tests, with Tukey HSD correction, were performed.

### Assessment of the overall tick vector competence for ASFV

Although our main assumption is that vector competence results are influenced by the experimental design, we attempted to assess the overall vector competence of the various *Ornithodoros* tick-ASFV associations. Based on the three main components described above, we scored each tick-ASFV association, according to the following ranking: “score 0” for ticks that were unable to become infected or for which it was impossible to conclude (IS = 0 or IS = NA, figure 1) ; “score 1” when ASFV was able to cross tick midgut and replicate in ticks; “score 2” when transmission from tick to pig was successful; and, “score 3” for ticks validating all components of vector competence, including ASFV transmission from ticks to other ticks. The attribution of a score was based on all data available from the different included references related to the considered tick-ASFV association. Attributing graduate scores of vector competence sounds as if that associated transmission events are also graduate. This is not the case: even if transmission to host scored 2 and sexual or transovarial transmissions scored 3, the success of tick-to-tick transmission does not necessarily imply the efficiency of transmission to host..

## Results

### Bibliometric description

The initial bibliographic search generated 236 references using Scopus and 138 by Pubmed, with 2 articles detected only by Pubmed. The ETHOS database of thesis manuscripts and our manual bibliography search allowed identifying 8 more resources. In this pool of bibliographic references, 39 references presenting vector competence original results were included in the final dataset. They consisted of 5 field studies monitoring natural tick infections, 34 articles of laboratory experiments on ticks and 2 mixed papers (Table 1). As shown in figure 2, the number of publications pertaining to the vector competence of *Ornithodoros* ticks for ASFV was consistently low over the years, while the total number of references related to ticks and ASFV progressively increased from the 1980s to 2022. The community of scientific experts identified is rather limited with only thirty-nine authors listed as first and last authors of the selected publications. After 2015, 9 out of 39 experts continued to publish in this domain and can be considered the current core of experts on the vector competence of ticks for ASFV.

**Figure 2.**
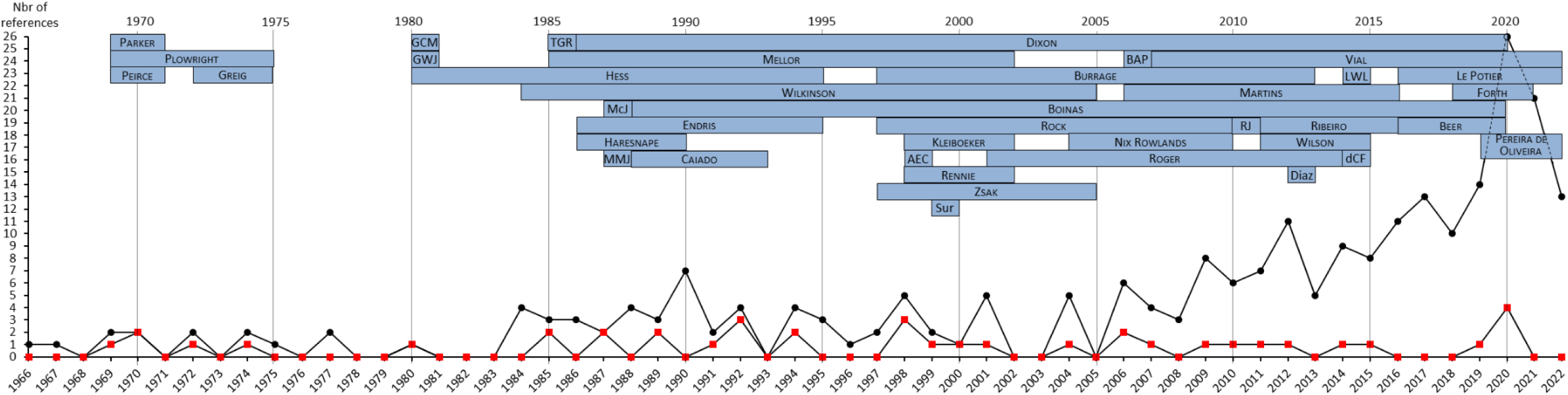
Number of bibliographic references published each year, from 1966 to 2022, on “African swine fever” AND “tick” (black dots = 246 screening references) and on the vector competence of *Ornithodoros* ticks for ASFV (red dots = 39 included references). The names of first and last authors of the 39 included references were reported in blue boxes at the top of the graphic. The length of boxes indicates their respective period of expertise, during which they published in the “African swine fever” AND “tick” fields (i.e., among the 246 screening references). List of the abbreviation author name: AEC: Anderson E.C.; BAP: Basto A.P.; dCF: de Carvalho Ferreira H.C.; GWJ: Gladney W.J.; GCM: Groocock C.M.; LWL: Loeffen W.L.; McJ: McCoy J.P.; MMJ: Monahan M.J.; RJ: Ravaomanana J.; TGR: Thomson G.R.

### Data related to the biological material: ticks and ASFV

A database was created to record all the extracted data. From the 39 included references, 2036 records corresponding to 900 unique experiments were extracted. For some references, authors detailed the results of their experiment for each tick or with multiple records over time for the same ticks. Each experiment was characterized by a combination of tick species, virus strain and experimental conditions (see Annexes 2 and 3). In total, the tick species and ASFV strains represented 51 tick-virus associations and more than half (34/51) appeared in only one publication.

For the ticks, ten species were tested for their vector competence for ASFV. The Iberian tick *O. erraticus* was the most studied (representing 44% of the reported experiments), followed by the *O. moubata* complex of species, including *O. porcinus* and *O. moubata* sensu stricto (43%). The North American *O. coriaceus* (6.4%) were third, then they were far followed by the other species *O. puertoricensis, O. verrucosus, O. savignyi, O. parkeri, O. sonrai, O. turicata* (3.7% - 1.0% - 0.8% - 0.4% - 0.4% - 0.2 % respectively). The geographical origins of ticks specified in the included references are presented in figure 3. All *O. erraticus* ticks tested came from Portugal, and most of them from the same region of Alentejo, while *O. moubata* and *O. porcinus* ticks came from diverse countries, which is representative of their large distribution range in East and Southern Africa.

**Figure 3.**
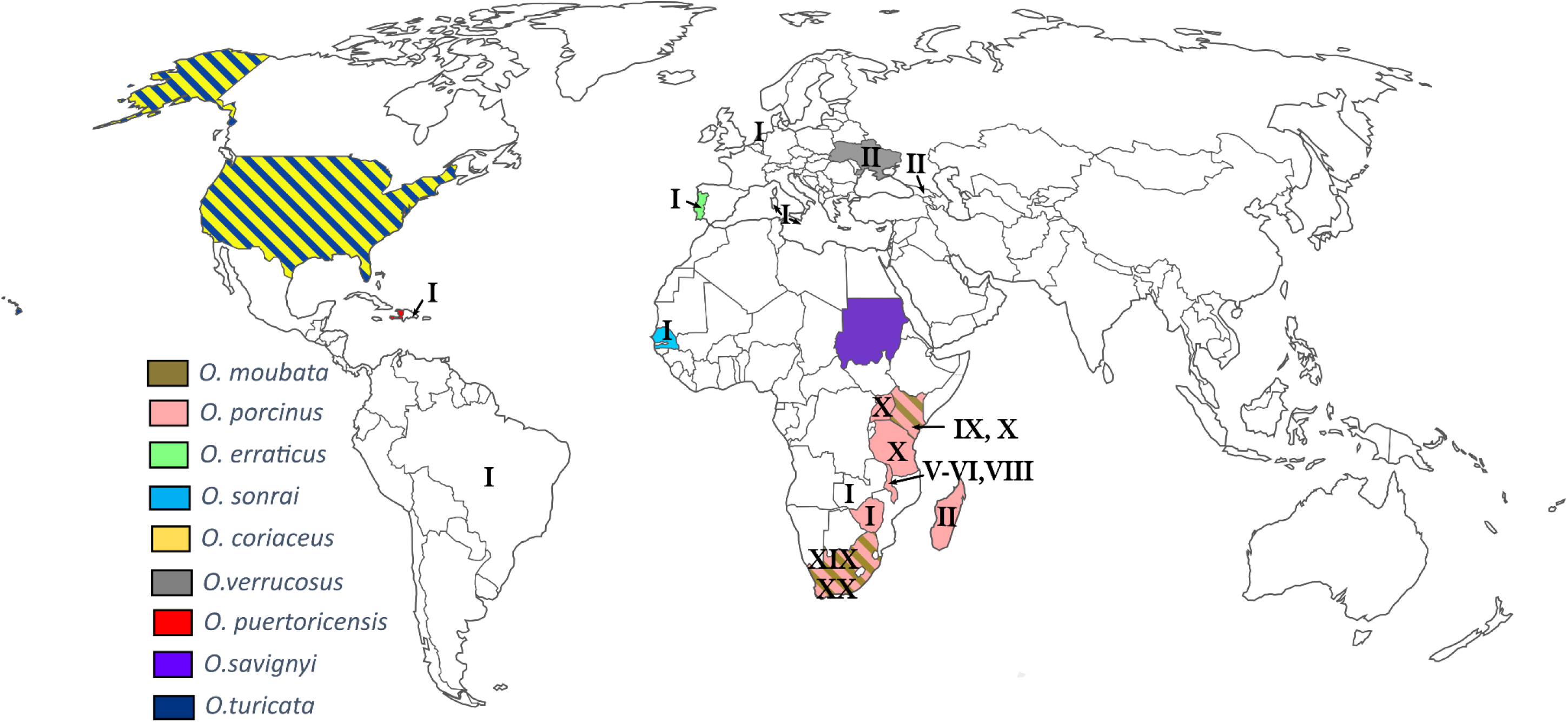
Countries of origin of *Ornithodoros* tick species (coloured according to the species) and ASFV genotypes (indicated by genotype number associated with each country), involved in tick-ASFV associations studied in included references. The origin of the tick *O. parkeri* was not indicated but should be the United-States, as described in the literature.

A total of 38 different ASFV strains were tested. Most ASFV strains used in the studies were isolated from ticks and pigs (43.1% and 40.4% of the reported experiments, respectively). Only four strains studied in 16.2% of the experiments were isolated from warthogs and 0.2% of virus strains are from undetermined hosts. ASFV strains were geographically distributed over 17 different countries, mostly in Africa, and belonged to 8 genotypes over the 23 genotypes described so far, with a large representation of the genotype I (64.9%, figure 3). The genotype of the 114a and Transvaal CV strains from South Africa remains unknown.

### Data related to the experimental design

Of the 900 experiments recorded, half used ticks directly sampled from the field (especially *O. erraticus* and *O. porcinus*) or reared in insectary for less than 5 years (48.7%). The other experiments used ticks reared in the laboratory for more than 5 years with some reared until 25 years for one study on *O. turicata* (47.2%). It was not possible to find information for the remaining experiments. The majority of experiments (57.9%) involved ticks in their later developmental stages, ranging from nymph 4 to adults. Early stages, from larva to nymph 3, were employed in 23.3% of the experiments. In the remaining experiments, authors either utilized a combination of early and late stages (14.7%) or did not provide information about the specific developmental stage used (4.1%). The number of ticks tested for each experiment was fairly low especially in laboratory experiments (median=7 ticks tested, mean=30.03, range=[1;1141]).

Four exposure routes were used to infect ticks. Natural blood feeding on hosts was reported either in field studies, or in some laboratory experiments using experimentally infected pigs. This method represented almost half of the total experiments conducted (47.4%). Two artificial feeding methods are reported: the longest established method uses capillary tubes inserted on tick mouthparts (8.8%) while a more recent approach uses animal skin, parafilm, or silicone membranes (35.7%). Another artificial exposure route was the syringe inoculation of ASFV directly in the tick haemocoel (8.1%).

ASFV infectious dose used to infect ticks was very heterogeneous and classified in three categories. Most experiments (78,4%) used medium to high viral doses (between 10^4^ and 10^6^ or over10^6^ HAD50/mL of blood). The DPE was very variable ranging from 0 day to up to 8 years. A DPE of 0 day is presumably used as a control for successful viral exposure. The longest reported DPE are from Boinas *et al*. (2011) and represent the duration between an ASFV outbreak and the date of virus detection in ticks in Portuguese farms, with virus detected in ticks up to 5 years after the outbreak. Nonetheless, more than half experiments (54.3%) reported a DPE lower than the duration required to differentiate a successful viral replication inside the tick from a detection of residual virus after the infection (61 days). Among them, one third performed virus detection in organs or transmission experiments, allowing conclusion on tick infection status.

### Factors influencing the likelihood of tick infection

On the initially 900 recorded experiments, we obtained a dataset of 407 observations with full information regarding tick infection status and experimental conditions (Annexes 4 and 5). Among the random effects tested, the tick species and virus genotype had no effect on the infection status but the viral strain had (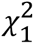=4.879 p-value= 0.027, see Annex 6 for details on viral strains). Model selection based on AIC ended up with three variables displaying large and comparable importance across all models: virus titre, tick stage and tick colony type (Figure 4 left). All those three variables had significant effects on infectious status (Figure 4 right).

**Figure 4.**
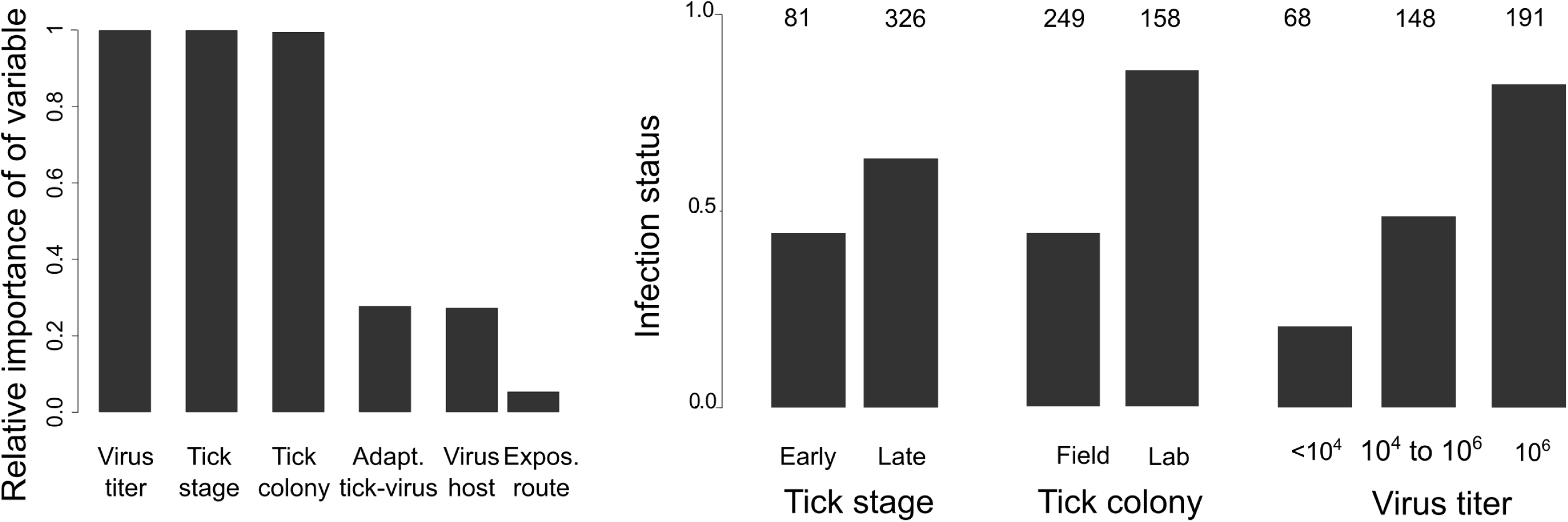
*Left*: Relative importance of explanatory variables across all models explaining infection status. The importance value for a particular variable is equal to the sum of the weights for the models in which the variable appears. “Virus host” is the host from which ASFV was primarily isolated. “Expos. route” is the exposure route for tick infection (natural, artificial, capillary, injection). *Right*: Effect of tick stage, tick colony and virus titre on the infection status (all three effects are significant). Mean of infection status (i.e., proportion of experiments where ticks were infected). Values above the bars are the number of experiments in each level. Virus titre in HAD50/mL of blood.

In more details, a higher viral titre corresponded to a higher likelihood of tick infection (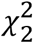= 38.493, p-value=4.423×10^−9^). Bigger nymphs and adults (late stages) were more prone to get infected than small nymphs (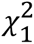=17.880, p-value=2.353×10^−5^). And ticks from laboratory colonies had higher probability of infection than ticks recently sampled in the field (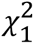= 14.031, p-value=1.798×10^−4^). The potential geographical co-occurrence of tick and virus, the exposure route for tick infection and the host from which ASFV was isolated were not significantly linked with infection status even if they were present in some of the best models (Annex 7). Using the dataset containing only experiments with adult ticks showed that sex had no influence on infection status.

### Vector competence scores for ASFV

Figure 5 summarizes the vector competence score for each of the 51 associations between soft ticks and ASFV virus tested so far. In more detail, 23.5% of the tick-ASFV associations presented a vector competence score null (score 0) i.e., ticks were unable to become infected or it was impossible to conclude on their infection status. For 27.5% of tick-ASFV associations, only tick infection (score 1) was achieved. Transmission from ticks to pigs was tested in 64.7% of tick-ASFV associations and was successful in 29.4% (score 2). Finally, 33.3% of tick-ASFV associations were assessed for ASFV transmission from ticks to ticks and 19.6% succeeded to transmit (score 3).

**Figure 5.**
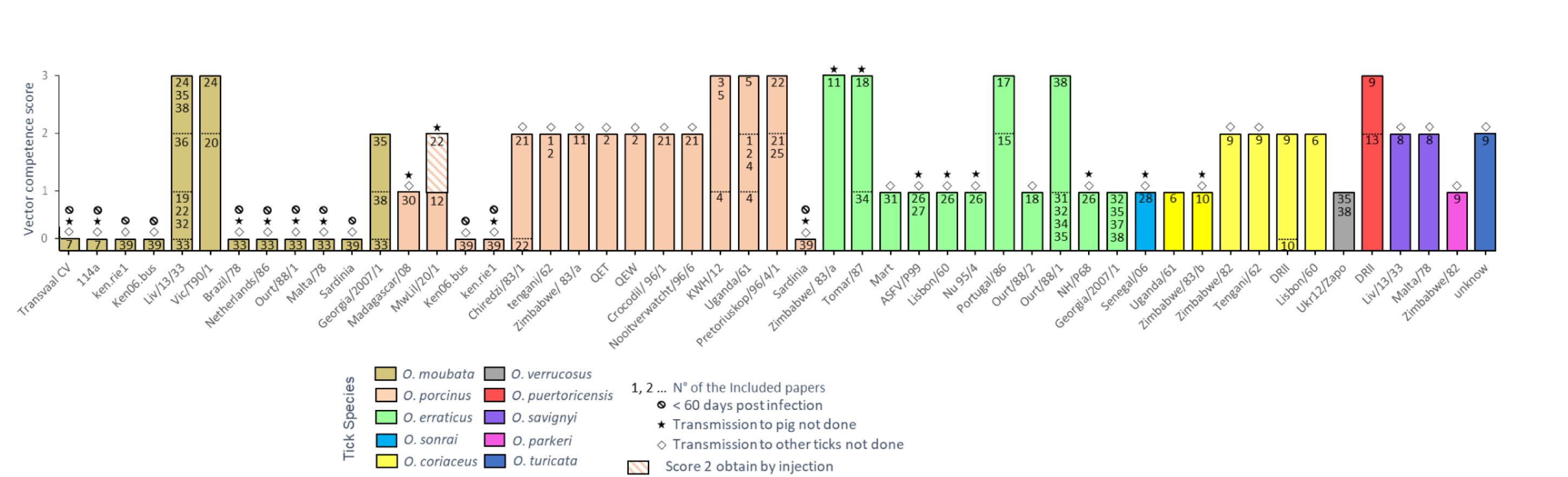
ASFV vector competence scores for the different tick-ASFV associations studied in the included references (score 0: no tick infection or no information to conclude, score 1: successful tick infection, score 2: successful ASFV transmission to pigs, score 3: successful ASFV transmission to other ticks). In each histogram bar, the numbers are related to the included references recorded in Table 1. For each included reference, an individual score is given according to available results in this paper. The overall vector competence score for a given tick-virus association is the best vector competence result reported from all included references related to this tick-ASFV association.

Only 27.5 % of tick-ASFV associations were fully tested for vector competence (including tick infection, transmission to pig, and transmission to other ticks) and 15.7% validated scores 1 to 3. Conversely, 3 associations reached score 3, without being tested for pig transmission (O. *erraticus*-Zimbabwe/83/a and O. *erraticus*-Tomar/87) or by failing to transmit to pigs when tested (O. *erraticus*-Ourt/88/1).

At the species level, most tick species tested were confirmed to be competent vectors (score 2) for at least one ASFV strain. This competence remains unclear for 3 species that were either tested with only one ASFV strain and failed to transmit the virus to pig and/or to ticks (*O. parkeri* and *O. verrucosus*) or never tested for transmission *(O. sonrai)*. Regarding ASFV transmission to other ticks, 6 tick species were tested and 4 (*O. moubata*, *O. porcinus, O. puertoricensis*. and *O. erraticus)* succeeded for at least one ASFV strain whereas 2 (*O. coriaceus* and *O. verrucosus)* failed. Finally, high heterogeneity of vector competence scores was observed within a same tick species depending on ASFV strains used (e.g. score 3 for *O. moubata*-LIV13/33 and score 0 for *O. moubata*-Brazil/78), and sometimes for a same tick-ASFV association (e.g. scores 0/1 and 2 for two *O. turicata* colonies of different ages with the same ASFV strain).

## Discussion

### Bibliographic resources

This paper is the first exhaustive review of all the original studies published on the vector competence of soft ticks for ASFV until 2024. Despite an exhaustive screening of bibliographic resources, only 39 references were considered and the expert panel had always been limited, as exemplified by only 9 publishing experts since 2015. This relative paucity of bibliographic resources regarding soft tick competence for ASFV may be attributed to the difficulty of investigating their vector competence, both in laboratory and in field conditions. Indeed, sampling endophilous *Ornithodoros* ticks and developing relevant methods to detect ASFV in natural tick populations (Jori et al., 2013; Oura et al., 2013) or rearing soft ticks and amplifying ASFV in laboratory conditions are long and laborious tasks (Vial, 2009; Carrascosa et al., 2011). In addition, although ASF is of high veterinary and economic concern, vectorial transmission is seen as occasional with a relatively minor role in disease spread at least in some regions (Guinat et al., 2016). Nevertheless, soft ticks play a key role in the maintenance of the virus and this is why characterizing competence for tick-virus associations is required to assess the risk of ASFV introduction and spillover, as Golnar *et al*. (2019) did for the US.

### Vector competence and tick-virus co-adaptation

Although all references included in this study aimed to evaluate the vector competence of soft ticks for ASFV, we noticed a considerable diversity of vector competence measurements that yield distinct information. As a consequence, vector competence was only partially assessed for most tick-ASFV associations, with a noticeable decrease of studies along the successive biological steps of vector competence. In addition, more than half of tick-ASFV associations were characterized by only one single article. Heterogeneous vector competence scores were observed within species between viral strains. This could be interpreted either as a lack of sufficient reliable studies or high tick-virus specificity.

Adaptation between arthropod vectors and viruses is based on genetic features defining susceptibility in both organisms and the selection of susceptibility may be mediated partly by detrimental effects of arboviruses on vector physiology (Gillespie et al., 2004). In the soft tick-ASFV model, the presence of ASFV MGF360 genes, the lectin-like EP153R gene, or the CD2-like EP402R gene enhance the entry and the replication of ASFV into ticks (Burrage et al., 2004; Rowlands et al., 2009). Some ASFV strains naturally lost these genes through prolonged circulation within the population of Iberian domestic pigs, or as a consequence of repeated passages on pig macrophages for vaccine production (Portugal et al., 2015). MGF 360 has also be shown to suppress the immune response of vertebrate and arthropod hosts against ASFV (Portugal et al., 2015). Arthropods, meanwhile, are known to display RNA interference as an important immune response against virus infection (Vijayendran et al., 2013). The involvement of such molecular pathway has recently been suggested in soft ticks, with ASFV-like integrated elements coding for small interfering RNA in *O. moubata* (Forth et al., 2020). More specifically, it appears that a greater number of these small RNA matched with ASFV genomes from genotype II than with genotype I, potentially contributing to the variation in vector competence between viral strains. This is in agreement with our results which showed significant variation in tick infection status depending on viral strain. However, no effect of virus genotype on tick infection status was found, which may be due to the very imbalanced dataset regarding virus genotype with genotype I being overrepresented compared to other genotypes.

We can suppose that field ticks, co-evolving or not with ASFV, may develop a much more heterogeneous and adaptive response against ASFV infection and show a lower success of infection, as it was already shown in some mosquito-virus models (Gillespie et al., 2004). Indeed, some authors previously reported very low ASFV infection rates in field ticks (Plowright et al., 1969; Haresnape & Wilkinson, 1989; Boinas et al., 2011). In the present study, laboratory tick colonies reared for more than five years were shown to be significantly much more infected by ASFV than field ticks or recently established laboratory ticks. In addition, ticks from old laboratory colonies may be more prone to efficient artificial blood feeding, thereby increasing the blood volume and the intake of ASF virus. However, the drawback of using old laboratory colonies is the risk of genetic drift and laboratory adaptation, leading to divergent phenotypes, compared to wild populations.

Despite heterogeneous vector competence according to studies and viral strain tested, experimental evidence shows that 7 *Ornithodoros* species from Africa (*O. moubata, O. porcinus and O. savignyi*), Europe (*O. erraticus*) and North America (*O. coriaceus, O. turicata and O. puertoricensis*) can be infected by at least one strain of ASFV and then transmit it to pigs.

### Effect of experimental factors on tick competence

Our analysis of vector competence references demonstrated the importance of the experimental design on vector competence results. The ASFV titre in the inoculum for infecting ticks appears as a key parameter in the vectorial competence experiments, with likelihood of tick infection above 50% when ASFV load was higher than 10^6^ HAD50/mL. Accordingly, a threshold of 10^5.75^ HAD50/mL was previously determined by Ribeiro (Ribeiro et al., 2015) for the specific ability of *O. erraticus* to maintain and multiply two Portuguese ASFV strains. Forth *et al*. (2020) also demonstrated an impact of ASFV titre on tick infection probability and load. Soon after ingestion, ASFV has to cross the tick midgut barrier to be amplified in replication units localized in tick endothelial cells and then enter the tick hemolymph (Burrage, 2013). This suggests that the tick midgut and other tick membranes act as porous surfaces; the probability of ASFV entry would thus depend on the number of available pores and on the virus load (Franz et al., 2015). Tick immune system such as RNAi could also block low level of ASFV infection (Forth et al., 2020).

The development stage of ticks had a significant impact on tick infection status, with better results for late stages (from nymph 4 to adults). This pattern confirms previous results on *O. erraticus*-Tomar and *O. erraticus*-OURT88/1 from Ribeiro (Ribeiro et al., 2015). This was explained by higher blood volume intake for late and large stages, and therefore highest ASFV titre inside the tick after blood meal digestion. In addition, even if female take a larger blood meal than males, we found no effect of observed sexual dimorphism in adults on the likelihood of tick infection which is in agreement with Ribeiro *et al*. (2015). In this study, the authors suggest that, in adults, the number of ASFV particles already reached a certain threshold necessary to cause tick infection.

The route for tick infection was expected to play a role on tick competence. Natural feeding on viraemic pigs may provide higher viral titre (usually >10^6^ HAD50/mL) than artificial feeding on membrane or with capillary (from 10^4^-10^6^ HAD50/mL) due to the difficulty for amplifying ASFV on pig macrophages. Despite differences between inoculation methods, this does not seem to translate into any significant differences on the success rate of infection in our dataset. This could be attributed to our dataset being incomplete and imbalanced, and with the virus titre being a confounding factor. Similarly, we did not find any effect of the initial host in which viral strain was isolated on tick competence.

## Conclusion

The examination of 39 references on the vector competence of *Ornithodoros* soft ticks for ASFV highlighted factors influencing vector competence results. For the same tick-ASFV association, vector competence results vary according to the experimental design employed. The selection of an experimental protocol constitutes a trade-off between scientific considerations and practical constraints. The findings of this meta-analysis provide a foundation upon which recommendations can be formulated for optimal protocols to ensure reliable results on vector competence for any soft tick-ASFV association. First, using late nymphs or adult ticks from a laboratory tick colony will improve the probability of obtaining engorged ticks, increase the volume of blood intake and allow testing for sexual or vertical transmission. Exposure should be conducted using a blood meal rather than by injection into the haemocoel. As the viral titre exerts the most significant influence on tick infection success, preferring blood from viraemic pigs or blood with a high titre will ensure sufficient virus load. Note that viraemic blood frozen at −80°C seems to result in efficient engorgement and infection rates (unpublished data). Once ticks are engorged, we recommend that viral DNA detection should not be used until 2 months after inoculation in order to differentiate between successful viral replication within the tick and detection of residual virus in the midgut. If viral DNA detection is performed on tick organs before 2 months post exposure, amplification of pig DNA on ASFV-positive samples is required to control for potential contamination by gut content that may have occurred during tick dissection (e.g. Bonsergent et al., 2023). Alternatively, the use of RT q-PCR with a reduced time frame is a viable option, as it detects ASFV gene expression. Such standardized procedures will facilitate further investigations on the vector competence of *Ornithodoros* ticks for ASFV, both expanding the range of tick species and virus genotypes tested and our general knowledge on the biological mechanisms involved.

## Author contributions

JB coordinated the study, collected all bibliographic resources, extracted the data. HJ-P, SM and JO analysed the data. LV and M-FLP coordinated the study. JB, HJ-P, LV, M-FLP and FB wrote the manuscript.

## Acknowledgements

The authors would like to thank Facundo Muñoz for his help with statistics.

## Data, scripts and codes availability

All data presented here and the script used for statistical analyses are available https://doi.org/10.18167/DVN1/RYLTJ7.

## Appendix

**Annex 1:**
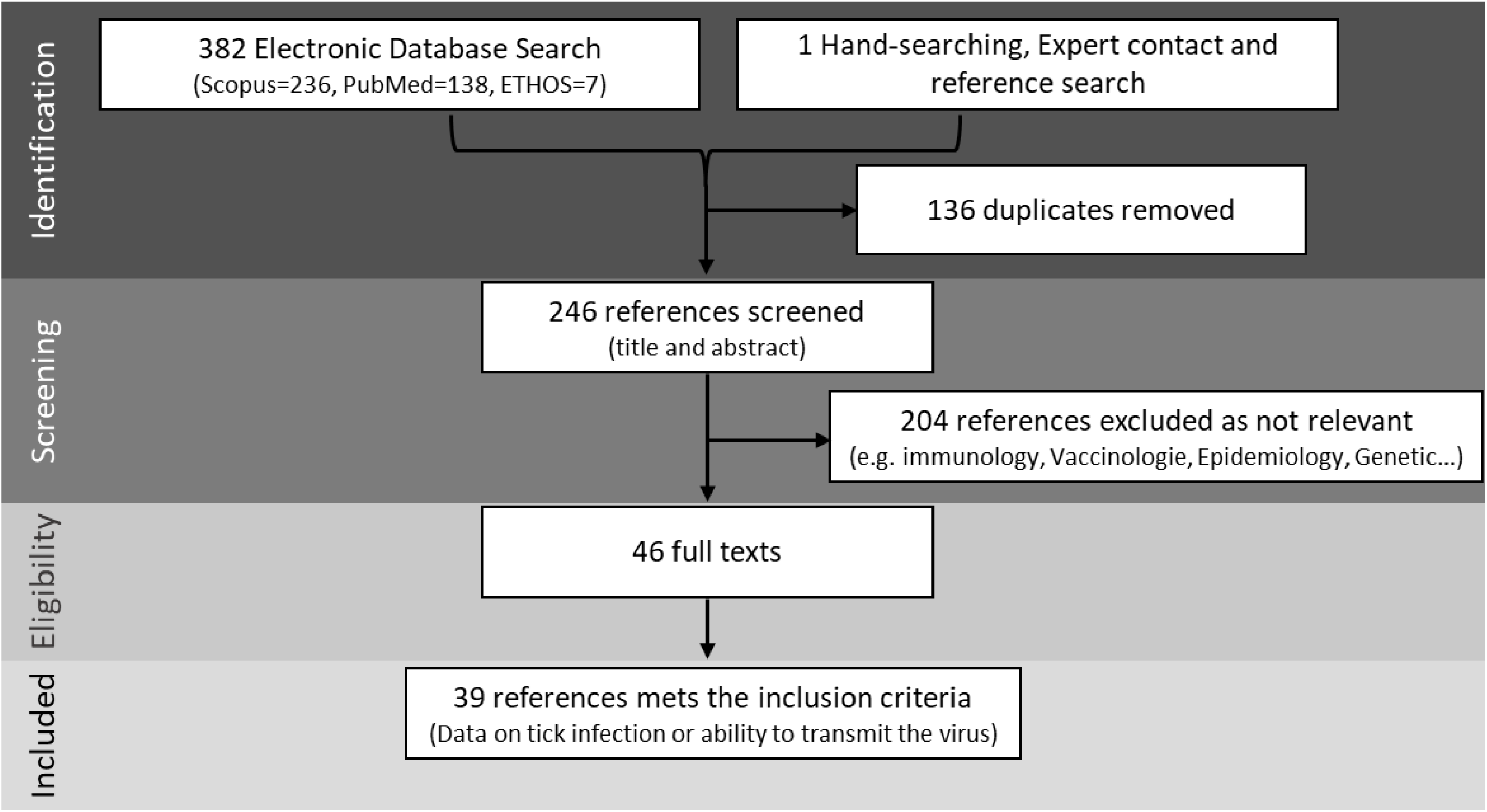
Article selection process according to the Preferred Reporting Items for Systematic Reviews and Meta-Analyses (PRISMA) guidelines. Except for 2 articles, all the articles found in PubMed were also found in Scopus, creating 136 duplicates that were removed. The larger number of references obtained in Scopus was expected since Scopus has larger scope and coverage than PubMed (AlRyalat et al., 2019)

**Annex 2:**
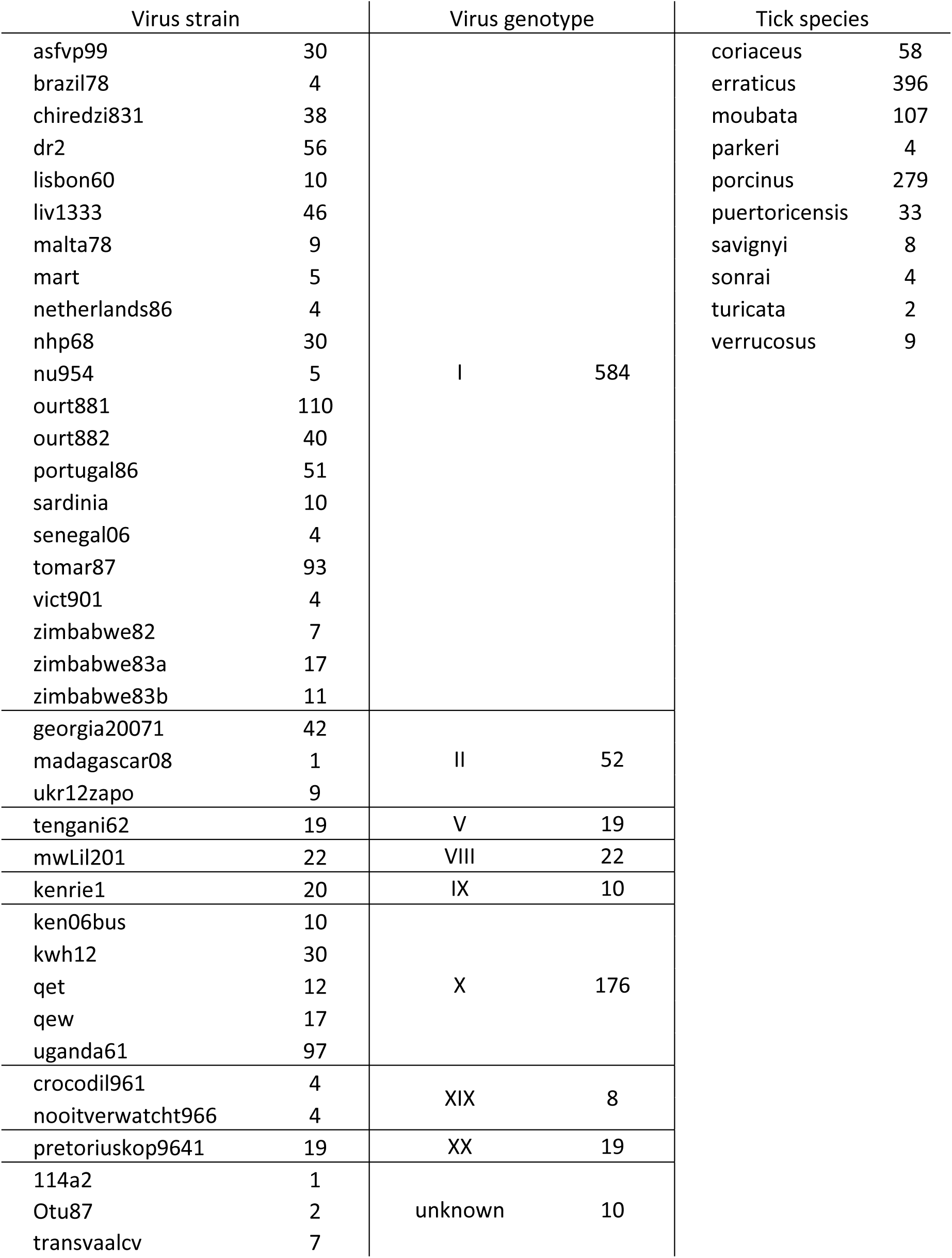
Number of experiments reported for each virus strain, virus genotype and tick species in the whole dataset (900 experiments)

| Virus strain |  | Virus genotype |  | Tick species |  |
| --- | --- | --- | --- | --- | --- |
| asfvp99 | 30 | I | 584 | coriaceus | 58 |
| brazil78 | 4 |  |  | erraticus | 396 |
| chiredzi831 | 38 |  |  | moubata | 107 |
| dr2 | 56 |  |  | parkeri | 4 |
| lisbon60 | 10 |  |  | porcinus | 279 |
| liv1333 | 46 |  |  | puertoricensis | 33 |
| malta78 | 9 |  |  | savignyi | 8 |
| mart | 5 |  |  | sonrai | 4 |
| netherlands86 | 4 |  |  | turicata | 2 |
| nhp68 | 30 |  |  | verrucosus | 9 |
| nu954 | 5 |  |  |  |  |
| ourt881 | 110 |  |  |  |  |
| ourt882 | 40 |  |  |  |  |
| portugal86 | 51 |  |  |  |  |
| sardinia | 10 |  |  |  |  |
| senegal06 | 4 |  |  |  |  |
| tomar87 | 93 |  |  |  |  |
| vict901 | 4 |  |  |  |  |
| zimbabwe82 | 7 |  |  |  |  |
| zimbabwe83a | 17 |  |  |  |  |
| zimbabwe83b | 11 |  |  |  |  |
| georgia20071 | 42 | II | 52 |  |  |
| madagascar08 | 1 |  |  |  |  |
| ukr12zapo | 9 |  |  |  |  |
| tengani62 | 19 | V | 19 |  |  |
| mwLil201 | 22 | VIII | 22 |  |  |
| kenrie1 | 20 | IX | 10 |  |  |
| ken06bus | 10 | X | 176 |  |  |
| kwh12 | 30 |  |  |  |  |
| qet | 12 |  |  |  |  |
| qew | 17 |  |  |  |  |
| uganda61 | 97 |  |  |  |  |
| crocodil961 | 4 | XIX | 8 |  |  |
| nooitverwatcht966 | 4 |  |  |  |  |
| pretoriuskop9641 | 19 | XX | 19 |  |  |
| 114a2 | 1 | unknown | 10 |  |  |
| Otu87 | 2 |  |  |  |  |
| transvaalcv | 7 |  |  |  |  |

**Annex 3:** Number of experiments reported in each category of the studied experimental factors in the whole dataset (900 experiments). NA: not available.

| Exposure route |  | Tick stage |  | Tick colony |  | Virus host |  | Viral titre |  | Geographic co-occurrence |  |
| --- | --- | --- | --- | --- | --- | --- | --- | --- | --- | --- | --- |
| artificial | 319 | early | 210 | field | 438 | pig | 364 | <10<sup>4</sup> | 109 | yes | 747 |
| capillary | 79 | late | 522 | lab | 425 | tick | 389 | 10<sup>4</sup> to 10<sup>6</sup> | 296 | no | 153 |
| natural | 427 | mix | 131 | NA's | 37 | warthog | 145 | >10<sup>6</sup> | 495 |  |  |
| inoculation | 75 | NA's | 37 |  |  | NA's | 2 |  |  |  |  |

**Annex 4:**
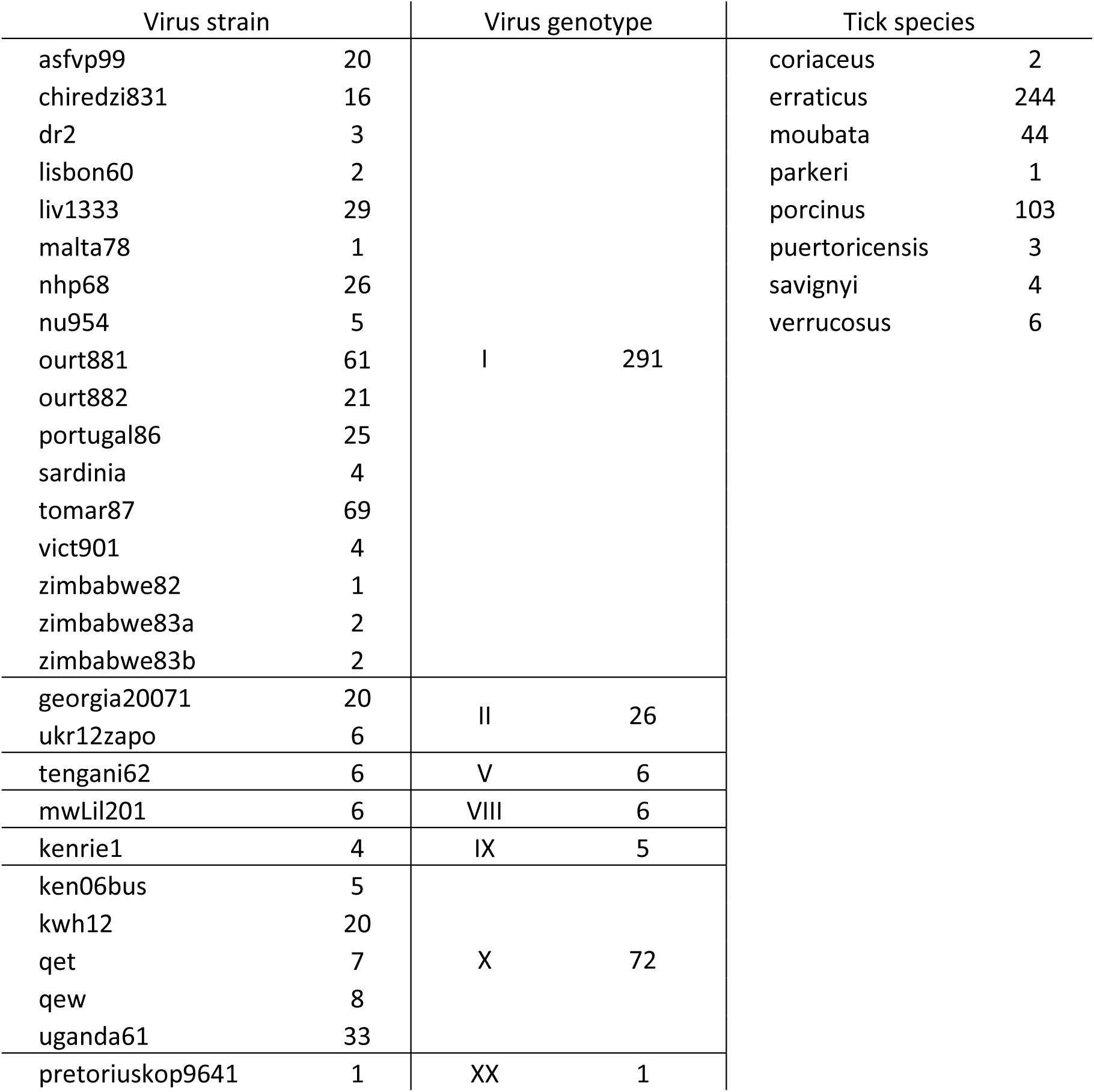
Number of experiments reported for each virus strain, virus genotype and tick species in the dataset analysed (407 experiments).

| Virus strain |  | Virus genotype |  | Tick species |  |
| --- | --- | --- | --- | --- | --- |
| asfvp99 | 20 | I | 291 | coriaceus | 2 |
| chiredzi831 | 16 |  |  | erraticus | 244 |
| dr2 | 3 |  |  | moubata | 44 |
| lisbon60 | 2 |  |  | parkeri | 1 |
| liv1333 | 29 |  |  | porcinus | 103 |
| malta78 | 1 |  |  | puertoricensis | 3 |
| nhp68 | 26 |  |  | savignyi | 4 |
| nu954 | 5 |  |  | verrucosus | 6 |
| ourt881 | 61 |  |  |  |  |
| ourt882 | 21 |  |  |  |  |
| portugal86 | 25 |  |  |  |  |
| sardinia | 4 |  |  |  |  |
| tomar87 | 69 |  |  |  |  |
| vict901 | 4 |  |  |  |  |
| zimbabwe82 | 1 |  |  |  |  |
| zimbabwe83a | 2 |  |  |  |  |
| zimbabwe83b | 2 |  |  |  |  |
| georgia20071 | 20 | II | 26 |  |  |
| ukr12zapo | 6 |  |  |  |  |
| tengani62 | 6 | V | 6 |  |  |
| mwLil201 | 6 | VIII | 6 |  |  |
| kenrie1 | 4 | IX | 5 |  |  |
| ken06bus | 5 | X | 72 |  |  |
| kwh12 | 20 |  |  |  |  |
| qet | 7 |  |  |  |  |
| qew | 8 |  |  |  |  |
| uganda61 | 33 |  |  |  |  |
| pretoriuskop9641 | 1 | XX | 1 |  |  |

**Annex 5:** Number of experiments reported in each category of the studied experimental factors in the dataset analysed (407 experiments).

| Exposure route |  | Tick stage |  | Tick colony |  | Virus host |  | Viral titre |  | Geographic co-occurrence |  |
| --- | --- | --- | --- | --- | --- | --- | --- | --- | --- | --- | --- |
| artificial | 169 | early | 81 | field | 249 | pig | 172 | <10<sup>4</sup> | 68 | yes | 369 |
| capillary | 46 | late | 326 | lab | 158 | tick | 174 | 10<sup>4</sup> to 10<sup>6</sup> | 148 | no | 38 |
| natural | 154 |  |  |  |  | warthog | 61 | >10<sup>6</sup> | 191 |  |  |
| inoculation | 38 |  |  |  |  |  |  |  |  |  |  |

**Annex 6:**
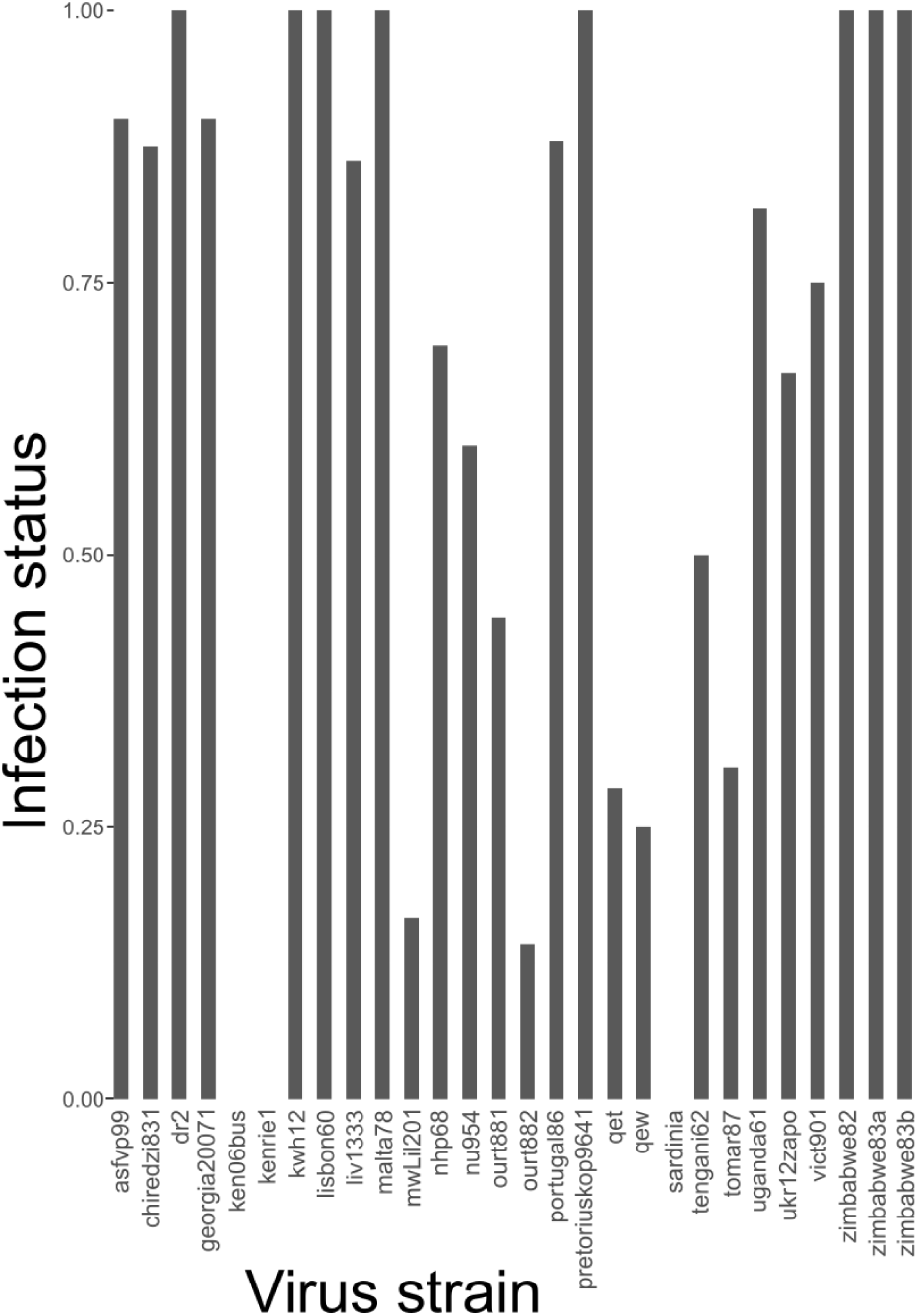
Mean tick infection status (i.e., proportion of experiments where ticks were infected) in relationship with virus strain tested.

**Annex 7:**
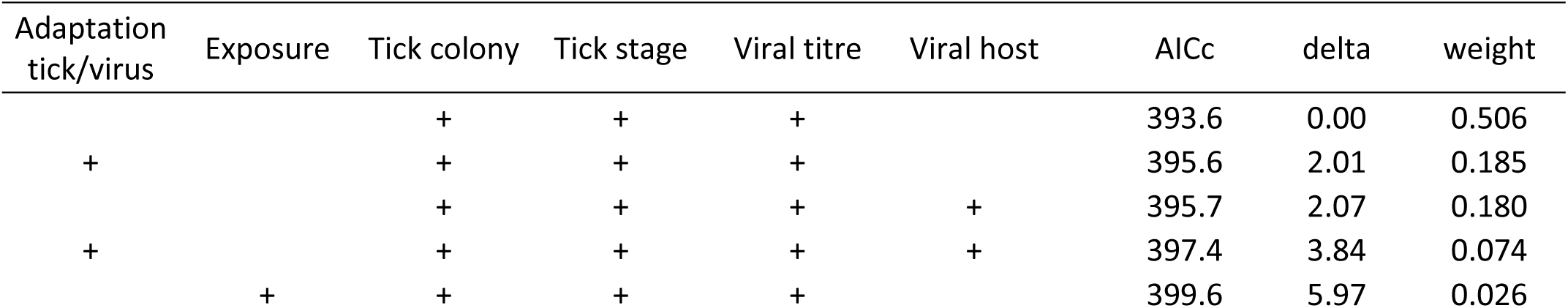
Five best models with the included variables influencing infection status (note that all models have «virus strain» as random effect), corrected Akaike Information Criteria and weight of the model (probability that the model is the best out of all models considered).

